# Genotype-Phenotype Correlation of T Cell Subtypes Reveals Senescent and Cytotoxic Genes in Alzheimer’s Disease

**DOI:** 10.1101/2021.10.19.464914

**Authors:** Dallin Dressman, Thomas Buttrick, Maria Cimpean, David Bennett, Vilas Menon, Elizabeth M. Bradshaw, Badri Vardarajan, Wassim Elyaman

**Author notes:** Correspondence should be addressed to: Wassim Elyaman Ph.D. or Badri Vardarajan Ph.D.

## Abstract

Recent studies identifying expression quantitative trait loci (eQTL) in immune cells have uncovered important links between disease risk alleles and gene expression trends in monocytes, T cells, and other cell types. However, these studies are generally done with young, healthy subjects, limiting the utility of their findings for age-related conditions such as Alzheimer’s disease (AD). We have performed RNA sequencing on four T cell subsets in genome-wide genotyped and well-characterized AD subjects and age- and sex-matched healthy controls from the Religious Orders Study/Memory and Aging Project. Correlating gene expression data with AD neuropathological traits, and with single nucleotide polymorphisms (SNPs) to detect eQTLs, we identified several significant genes involved in T cell senescence and cytotoxicity, consistent with T cell RNA sequencing studies in aged/AD cohorts. We identified unexpected eQTLs previously associated with neuropsychiatric disease traits. Finally, we discovered that pathways related to axon guidance and synaptic function were enriched among trans-eQTLs in coding regions of the genome. Overall, our data sheds more light on the genetic basis behind phenotypic changes in T cells during aging and AD.

## Introduction

Alzheimer’s disease (AD) is a progressive and incurable neurodegenerative disease of aging. While the early-onset form of AD is familial, involving rare mutations in the amyloid precursor protein (*APP*) and/or the presenilins, over 90% of AD cases are late-onset and sporadic (Bekris et al. 2011). Early neuropathological findings consistently identified extracellular plaque, composed of the amyloid beta fragment of APP, and neurofibrillary tangles, composed of tau protein aggregates, in post-mortem AD brain samples (Price et al. 1991). More recently, genetic and other studies have highlighted markers of inflammation and immunity as important components of AD progression. Microglia, the resident immune cells of the central nervous system (CNS), contribute to many important homeostatic processes in the brain and express a handful of key AD risk genes (Sims et al. 2017, Olah et al. 2018).

While microglia are currently thought to accomplish most of the immune surveillance in the healthy brain, several neurological conditions are known to involve an influx of peripheral immune cell types into the CNS, including blood-derived monocytes and T cells (Kramer et al. 2019; Brochard et al. 2008; Varvel et al. 2016). An immunohistochemical study hinted at a correlation between T cell influx and tau pathology in AD, suggesting that this phenomenon may occur in mid- to late-stage neurodegeneration (Merlini et al., 2018). Recently, using single-cell sequencing technology, we have identified a population of cytotoxic T cells in post-mortem and surgically resected brain tissue from individuals with epilepsy or AD; many of these T cells express markers of late differentiation and senescence (Olah et al. 2020). Others have found similar results in cerebrospinal fluid of AD patients (Gate et al. 2020). A number of researchers have employed various methods to modulate T cell populations in mouse models of AD. Results from these studies often vary based on the model used, disease stage, T cell subtype, or T cell antigen specificity; however, these studies implicate T cells in disease regulation and pathogenesis (Cao et al. 2009; Baruch et al. 2015; Mittal et al. 2019).

Effector T cells are generally classified as either cytotoxic (CD8+) or “helper” (CD4+) T cells. CD8+ T cells can directly kill target cells, usually by Fas-FasL signaling, release of cytotoxic enzymes or proinflammatory enzymes, or a combination of these (Farhood, Najafi & Mortezaee 2018). CD8+ T cells employ these tactics upon encountering intracellular antigens presented by the major histocompatibility complex (MHC) class I, usually in the context of viral infection or tumor growth (Wherry & Ahmed 2004; Ahmadzadeh et al. 2009). CD4+ T cells, on the other hand, generally play a more supportive role, participating in cytokine signaling to stimulate other adaptive or innate immune cells (Zhu et al. 2010). Common subtypes of CD4+ T cells are traditionally delineated by the cytokines they release (Dressman and Elyaman 2021). T helper (Th)1 release pro-inflammatory cytokines like interferon (IFNγ) or tumor necrosis factor (TNFα), Th2 produce generally anti-inflammatory cytokines such as interleukin (IL)-4 and IL-13, Th17 is known for IL-17 release, and regulatory T cells (Tregs) secrete cytokines with a more immunosuppressive function, including IL-10 and TGF-β (Luckheeram et al. 2012, Zhu et al. 2010). T cells are considered “naïve” before encountering their specific antigen. Once primed with an antigen, T cells divide and differentiate, and long-lived memory T cells are developed with the capacity to expand rapidly in response to the same antigen (Sallusto et al. 2003). Naïve T cells have, until recently, been considered synonymous with CD45RA-expressing T cells, and memory T cells with CD45RO+ cells. However, a subset of memory T cells, termed TEMRA, is now known to lose CD45RO expression and re-express CD45RA (Tian et al. 2017).

Thus far, few groups have explored possible genetic underpinnings of T cell phenotypic changes in aging and AD. Several expression quantitative trait loci (eQTL) studies in AD patients use brain tissue samples (GTEx Consortium 2017, Ng et al. 2017), providing helpful spatial context, but cannot be utilized to describe cell type-specific changes. Raj et al. (2014) investigated eQTLs in monocytes and naïve CD4+ T cells; and while their analysis focused on AD genetic variants, only healthy young subjects were used in the study, and any T cell subtype-specific effects would escape detection. In this study, we aim to better characterize the genetic risk factors behind T cell gene expression changes in AD subjects and controls. We conducted RNA sequencing of both CD4+ and CD8+ T cells sorted from peripheral blood mononuclear cells from patients in the ROSMAP cohort (Bennett et al. 2018). We further divide the two main T cell subsets based on the expression of CD45RO, resulting in the following 4 subsets: CD4+CD45RO-, CD4+CD45RO+, CD8+CD45RO-, and CD8+CD45RO+ T cells. Using this sequencing data, we identified three gene co-expression modules, and correlated gene expression with AD pathological traits and with whole genome sequences to detect eQTLs. We discovered that several genes commonly associated with cytotoxicity and senescence were found in association with AD neuropathological traits, in co-expression modules, and, unexpectedly, in eQTLs associated with neuropsychiatric disease.

## Results

### Comparison of Gene Expression Between T Cell Subsets and AD Neuropathological Traits

To better understand the phenotypic characteristics of CD4+ and CD8+ T cells in AD subjects, we leveraged cryopreserved peripheral blood mononuclear cells (PBMCs) from 48 AD patients and 48 age- and sex-matched control subjects in the ROSMAP cohort (**Figure 1a**). We used flow cytometry to sort live naïve and memory CD4+ and CD8+ T cells based on the expression of CD45RO. Briefly, 96 cryopreserved PBMC samples were thawed and were stained with the following surface markers: CD3, CD4, CD8, and CD45RO. Four T cell subsets were sorted from each subject: CD3+CD4+CD45RO-, CD3+CD4+CD45RO+, CD3+CD8+CD45RO-, and CD3+CD8+CD45RO+ T cells. Cells were lysed to collect RNA for 3’-digital gene expression (3’-DGE) sequencing (see Methods, and **Supplementary Table 1** for raw data). We compared gene expression between the four T cell subsets, and between AD and control subjects within each subset. Surprisingly, very few genes were differentially expressed when comparing AD and age- and sex-matched healthy control subjects within a given T cell subset (**Figure 1b)**. Most of the significant differences in gene expression were found in comparisons between T cell subsets, especially comparing CD4+ to CD8+, regardless of AD status.

**Table 1:** Total numbers of cis- and trans-eQTL, including unique SNPs and genes, among the four T cell subsets studied.

| T cell subset | # of cis-eQTL | Cis-eQTL unique SNPs | Cis-eQTL unique genes | # of trans-eQTL | Trans-eQTL unique SNPs | Trans-eQTL unique genes |
| --- | --- | --- | --- | --- | --- | --- |
| CD4+CD45RO- | 37 | 32 | 23 | 43721 | 16330 | 5234 |
| CD4+CD45RO+ | 14 | 14 | 14 | 32550 | 12246 | 5224 |
| CD8+CD45RO- | 5 | 5 | 5 | 26315 | 11293 | 4566 |
| CD8+CD45RO+ | 32 | 31 | 23 | 64281 | 19318 | 5090 |

**Figure 1:**
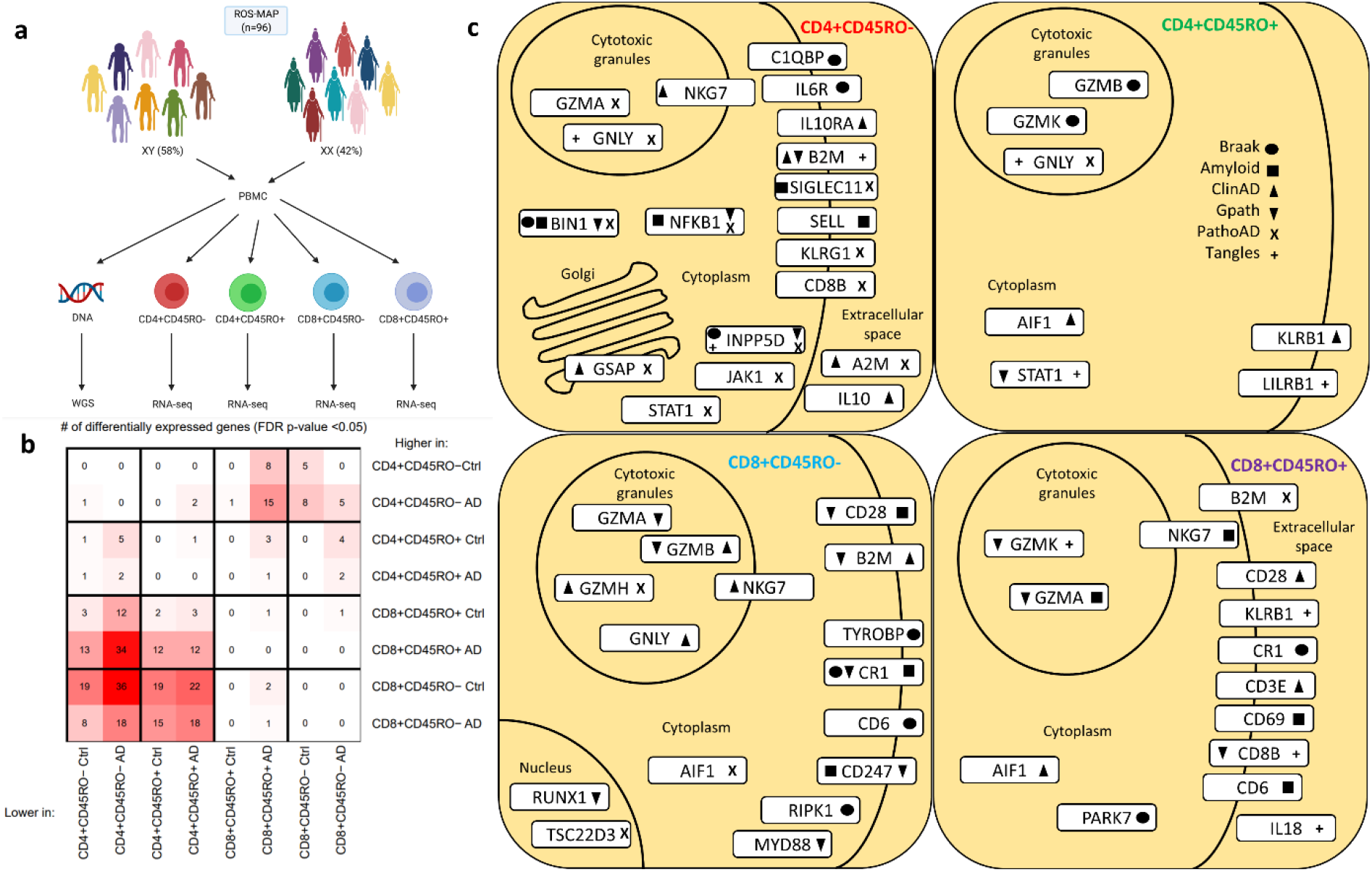
Patterns of T cell gene expression, co-expression, and correlation with AD pathological traits. **a**) Overview of study design, including number of AD and control subjects, cell sorting strategy, and data collection. Created with Biorender.com. **b**) Heatmap displaying numbers of differentially expressed genes (DEGs) when compared between given cell types and AD status. n = 96, FDR-adjusted p-value < 0.05. **c**) Selected genes (with immune or disease significance) associated with AD pathological traits (shape legend shown at top right) at a nominal *p* < 0.05 for each cell type, arranged according to the cellular localization of protein product.

To derive further meaning from gene expression data, we tested the correlation of each gene’s expression with different neuropathological measurements of AD severity from ROSMAP. These include Braak staging of tau tangle pathology, pathological or clinical diagnosis of AD (PathoAD/ClinAD), global AD pathology burden (Gpath), and the presence of amyloid beta plaque or tau tangles. Interestingly, several genes involved in cytotoxicity, inflammation, and T cell activation arose (**Figure 1c**). We identified hundreds of genes in all four T cell subsets that have a nominal relationship with one or more AD neuropathological traits (for a full list, see **Supplementary Table 2**). Of note, several granzyme genes were common in these associations, as well as senescence-related genes such as *GNLY* and *NKG7*, AD risk genes such as *CR1* and *BIN1* (Lambert et al. 2009, Wijsman et al. 2011), and markers of inflammation and T cell activation such as *NFKB1* and *CD28* (Bank et al. 2014, June et al. 2009). While the role played by senescent or pro-inflammatory T cells in pathological mechanisms of AD is only beginning to come to light, other studies have shown an upregulation of several of these senescence and late activation markers in T cells from aged individuals and AD patients (Hashimoto et al. 2019; Gate et al. 2020; Xu and Jia 2021).

### Gene Co-Expression Network Analysis

We also sought to detect broader trends of gene expression and co-expression across subjects. To this end, we used weighted gene co-expression network analysis (WGCNA) to detect co-expression modules whose collective expression differed consistently across samples. We found three such modules, the first of which was most highly expressed in memory CD8+CD45RO+ T cells, and the third most upregulated in naïve CD8+CD45RO- T cells (**Figure 2a)**. Module 1 featured several genes associated with IL-10 signaling, including *TSC22D3*, an anti-inflammatory transcription factor that is known to be stimulated by glucocorticoids and interleukin 10 (Berrebi et al. 2003), that we previously found to be highly expressed in a cluster of CD8+ T cells in AD and epilepsy (Olah et al. 2020). Module 2 included multiple genes associated with ribosomal function, suggesting an overall association with protein synthesis. Module 3 contained several genes often associated with senescence and cytotoxicity, including *KLRF1, KLRG1, GZMA, GZMB, GZMH, GZMK, CST7, GNLY*, and *NKG7* (**Supplementary Table 3**, Gothert et al. 2012, Pappalardo et al. 2020, Susanto et al. 2012, Wang et al. 2013). Interestingly, module 1 and module 3 genes were among those detected in association with AD neuropathological traits (**Figure 1c**).

**Figure 2:**
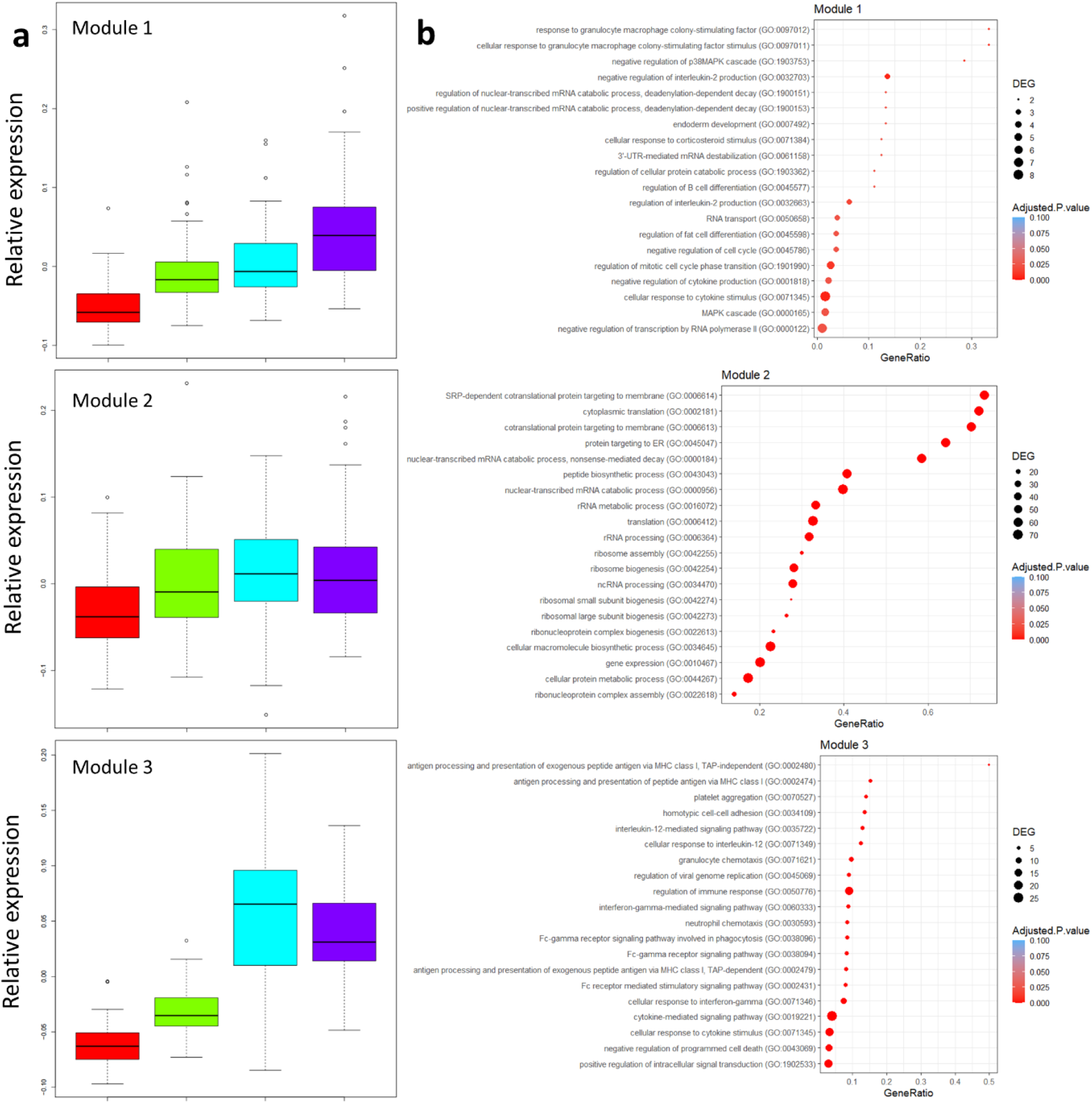
Relative abundance of WGCNA co-expression modules by cell type and pathway analysis. **a**) Relative expression of modules 1, 2, and 3 are shown for four T cell subtypes (red = CD4+CD45RO-, green = CD4+CD45RO+, cyan = CD8+CD45RO-, magenta = CD8+CD45RO+). **b)** Results of Enrichr pathway analysis on modules 1, 2, and 3. Dot size reflects the number of module genes in a given pathway, location on the X axis represents the proportion of the GO pathway represented by module genes, dot color represents p-value. Terms on the left are from the GO Biological Processes 2021 database.

We conducted unranked pathway analysis using Enrichr on module genes with GO Biological Processes as the input database. Upregulated pathways among module 1 genes include responses to cytokines such as IL-2 and GM-CSF (**Figure 2b**), while module 3 involved pathways related to IL-12 and IFN-γ signaling, as well as antigen processing. IL-12 is classically understood to promote the differentiation of Th1 type CD4+ T cells (Vacaflores et al. 2016), while IFN-γ is a potent part of the antiviral immune response (Ruby & Ramshaw 1991). These findings suggest that T cells in aged individuals exhibit a senescent pro-inflammatory phenotype that may be implicated in viral clearance.

### Expression Quantitative Trait Loci

Recent studies have advanced our understanding of the genetic underpinnings of AD pathology by detecting links between genetic variants and quantitative traits such as MRI imaging metrics (Shen et al. 2010), gene expression from various cell types (Raj et al. 2014, Chen et al. 2016, Kasela et al. 2017), or protein levels in brain, CSF, and plasma (Yang et al. 2021). To better elucidate the genetic basis of T cell phenotypic changes in AD, we correlated gene expression with genomic data previously generated from ROSMAP participants to identify eQTLs. We first tested for cis-eQTL associations, involving single nucleotide polymorphisms (SNPs) within 500 kb of a gene’s coding region. We defined significant cis-eQTLs as those with a false discovery rate under 0.05, whose gene was expressed in at least 20% of individuals with a maximum expression count of 3 or more (for a full listing of cis-eQTLs in all T cell subtypes, see **Supplementary Table 4**). Trans-eQTLs were defined by the SNP and gene coding region at least 500 kb apart, or on different chromosomes, and significance at FDR<0.05. Tens of thousands of significant trans-eQTLs were detected in each T cell subset (full listing in **Supplementary Table 5)**, with over 10,000 unique SNPs and ~5,000 unique genes in each subset (**Table 1**).

Most genes whose expression was correlated with trans-eQTLs were shared across all T cell subsets, while the SNPs themselves were subset-specific in their association with genes (**Figure 3a**). We visualized the distribution of trans-eQTL across the genome by generating Manhattan plots to summarize the location of SNPs found in trans-eQTL by chromosome (**Figure 3b)**.

**Figure 3:**
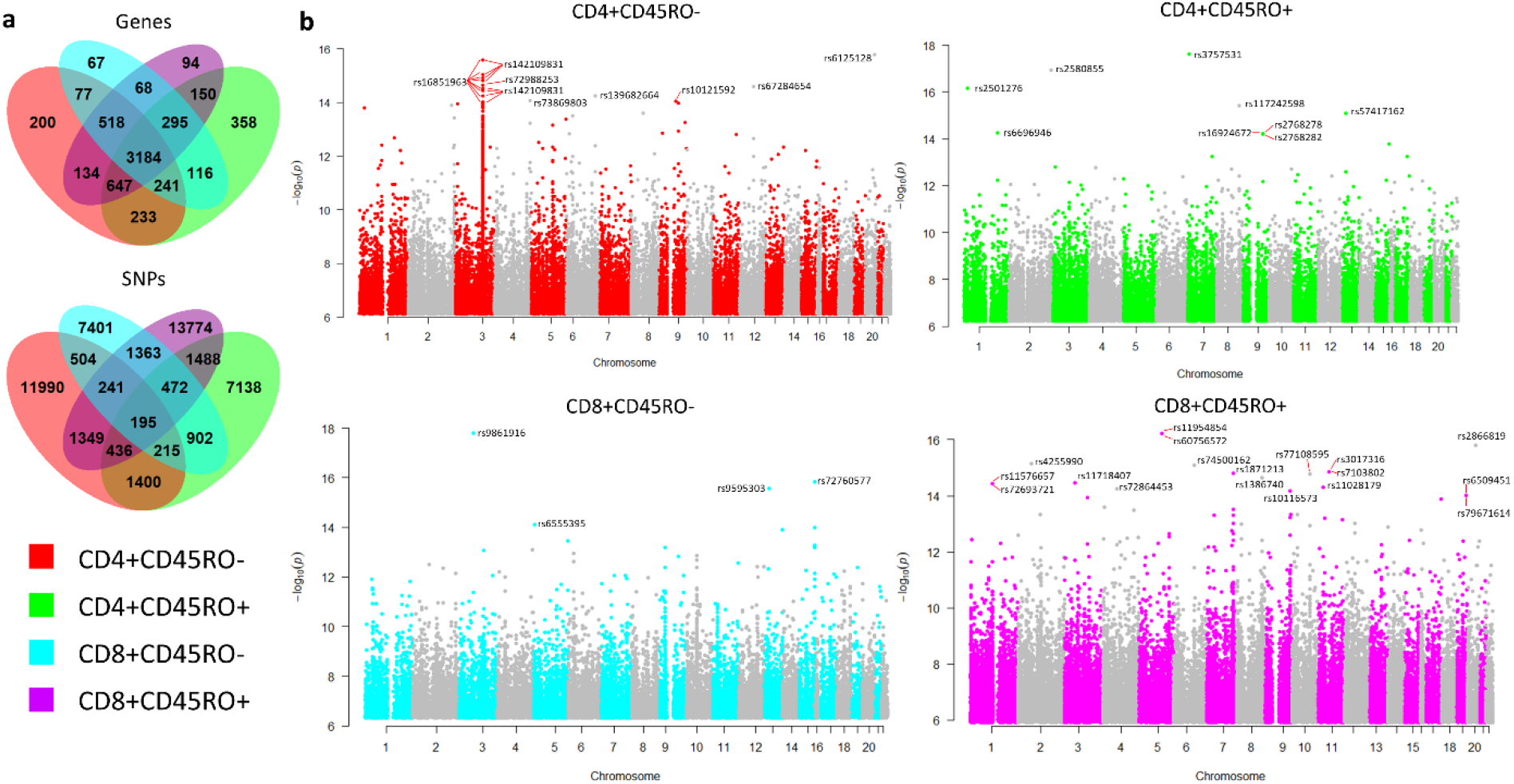
Distribution of trans-eQTL across the genome and between T cell subtypes. **a**) Venn diagrams showing distribution of unique genes (top) and SNPs (bottom) among the four T cell subsets. Genes found in trans-eQTL are mainly shared among all T cells, while SNPs are more subtype-specific. **b**) Manhattan plots showing the distribution of SNPs from the trans-eQTL in each cell type, with SNPs at –log_10_(*p*) > 14 annotated.

Owing to the age of our cohort, we recognized that gene expression changes in the T cell subsets may be driven by variants associated with neurological or immunological conditions other than AD. To assess disease association, we compared trans e-QTLs to SNPs in the GWAS catalog database (see **Supplementary Table 6)**, restricting to traits in four disease categories (see Methods and **Supplementary Table 7** for categorization of GWAS traits). Several dozen SNPs among the trans-eQTLs have been shown to associate with traits related to immune function, neuropsychiatric conditions, autoimmune disease, or neurodegenerative disease, in genome-wide association studies (**Figure 4a**). Most of these SNPs were trans-eQTLs from CD4+CD45RO+ cells, with neuropsychiatric conditions featuring prominently among the disease-associated SNPs in this cell type. Circos plots display the distribution of these GWAS catalog-associated trans-eQTLs across the genome (**Figure 4b**). Interestingly, among the disease-associated eQTLs in CD4+CD45RO+ cells were several immunosenescence and cytotoxicity-related genes, including *GNLY*, *GZMH*, *NKG7*, and *PRF1*, all of which were regulated by SNPs associated with anxiety.

**Figure 4:**
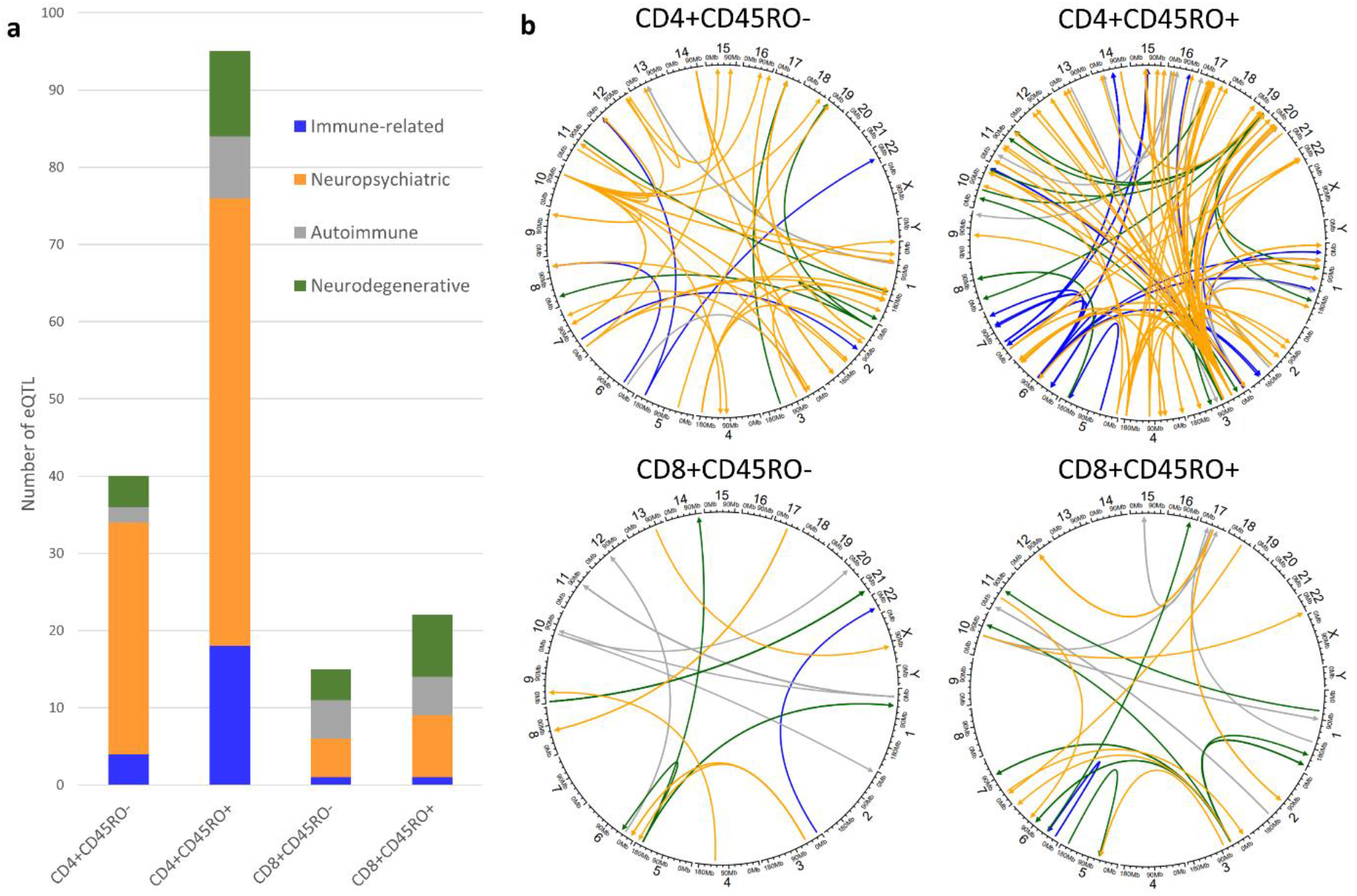
Disease association of trans-eQTL. **a**) Stacked bar graph showing numbers of trans-eQTL whose SNP is found to associate with certain categories of disease traits as found in the GWAS catalog. **b**) Circos plots displaying the distribution of disease-associated trans-eQTL by chromosome. Color legend for GWAS categories is found on the bar graph.

Finally, we sought to highlight overall functional trends in both the SNPs and the differentially expressed genes of the trans-eQTLs. In this way, we could better describe the mechanisms behind regulation of T cell gene expression in AD, as well as the nature of phenotypic changes specific to each T cell subset. To elucidate functional and mechanistic processes in trans-eQTLs, we conducted pathway enrichment analysis. For SNPs from the trans-eQTL, we first determined SNPs found within a gene (between the transcription start site and end site), then ran unranked pathway analysis on those genes with Enrichr. Unexpectedly, these genes were overwhelmingly found in pathways related to neuronal development, axon guidance, and synaptic transmission in all four T cell subtypes (**Figure 5**). Pathway analysis of genes from the trans-eQTL returned GO terms related to general RNA processing and translation in all four T cell subsets (**Supplementary Figure 1**).

**Figure 5:**
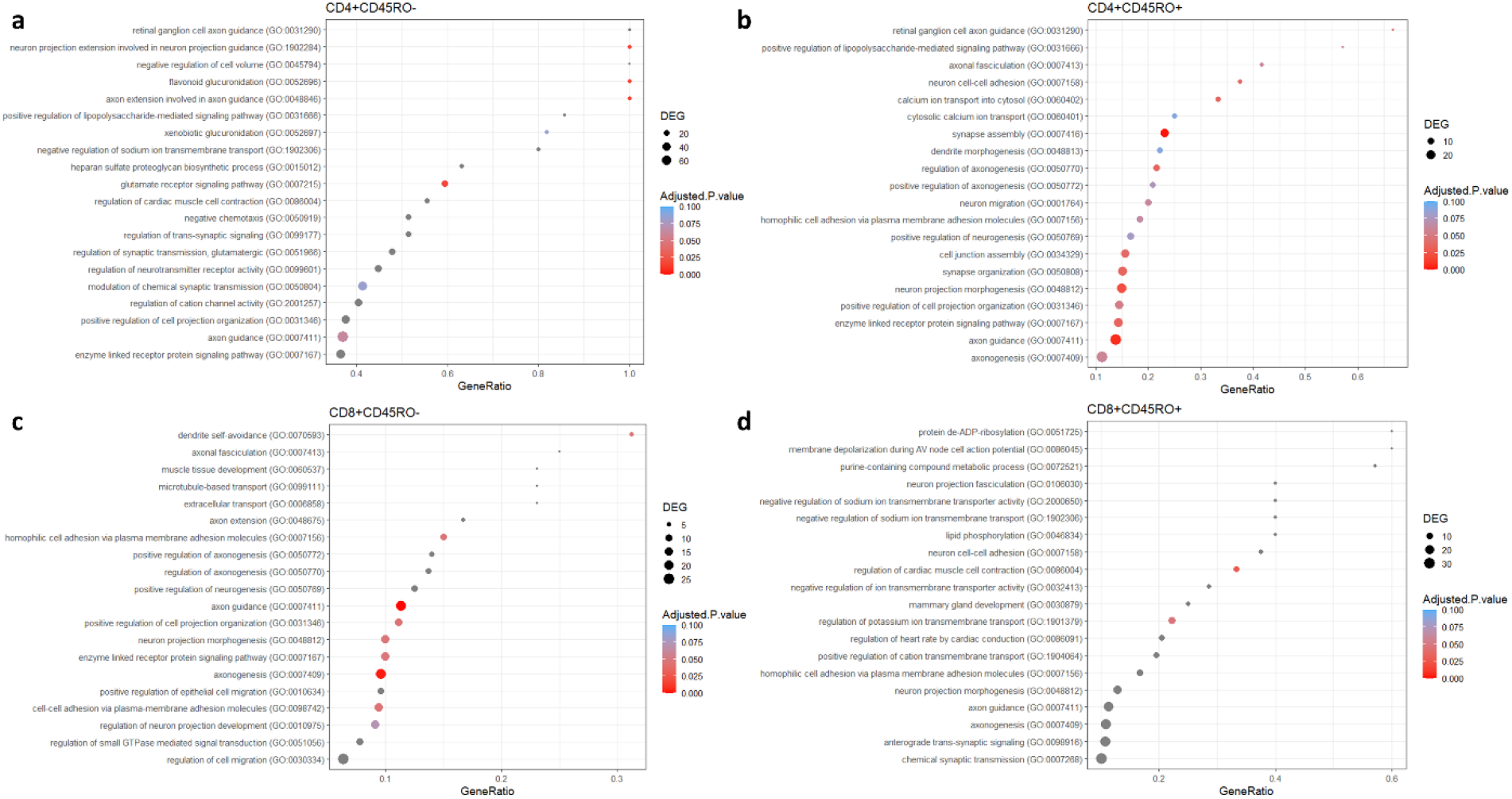
Pathway analysis of trans-eSNPs within the coding region of a gene. For SNPs that fell within the coding region of a gene, pathway analysis was run with Enrichr against the GO Biological Processes 2021 database for each cell type, **a**) CD4+CD45RO-, **b**) CD4+CD45RO+, **c**) CD8+CD45RO-, **d**) CD8+CD45RO+. These SNPs were highly associated with pathways related to synapse organization and function, axon guidance, and neurodevelopment.

## Discussion

This study is the first, to our knowledge, to examine the genetic underpinnings of T cell expression phenotypes in aged individuals and AD patients. Our use of CD45RO marker in cell sorting yields a preliminary look into how naïve and memory T cell phenotypes may differ from each other in an AD context. Furthermore, the neuropathological data collected from the ROSMAP cohorts allows us to correlate T cell expression data directly with AD-relevant traits. These findings allow us to consider the genetic basis for changes in T cell behavior during AD, and how T cell function and CNS AD pathology may influence each other.

From the expression data alone, our co-expression analyses highlight genes with key functional relevance, which may contribute to the neurodegenerative process. Module 3 is especially enriched in genes with direct cytotoxic function, such as granzyme genes, and others that are upregulated in T cells that have become senescent and pro-inflammatory, such as *GNLY* and several *KLR* family members. Module 1, conversely, includes several genes that participate in IL-10 signaling, which is classically understood to be anti-inflammatory, especially for T cells. Interestingly, module 3 is most highly associated with the CD8+CD45RO- subset, suggesting a more naïve phenotype for these cytotoxic and senescent T cells, although this expression profile may arise from the late-differentiation TEMRA phenotype as well, which is also CD45RO-. Several module 1 and module 3 genes also appear in association with AD neuropathological traits, as do some risk genes for AD and Parkinson’s disease (Fig. 1b), although these are only nominally significant.

Our eQTL analyses resulted in tens of thousands of trans-eQTL for all four T cell subsets, despite a relatively small sample size. The trans-eQTL involved over 10,000 unique SNPs and 5,000 unique genes in each T cell subtype. Existing studies of eQTL in T cells often detect SNPs that are risk variants for autoimmune disease, as might be expected. Strikingly, after correlating SNPs among our trans-eQTL with disease-associated SNPs, we found many that matched traits related to neuropsychiatric conditions such as schizophrenia and depression. Because few studies have examined potential connections between T cell phenotypes and psychiatric disorders (Pape et al. 2019), these associations may warrant a further look into how T cells may be altered, or alter nervous system function, in these conditions.

When we determined where SNPs were located among the T cell trans-eQTL, we were surprised to find that SNPs within gene coding regions were often found within genes related to axon guidance and neuronal process growth, as highlighted in our pathway analysis results. While it is important to focus on how T cells might drive neuronal death and dysfunction in conditions like AD, these findings suggest that genes that shape nerve growth during development and throughout life may also serve to regulate T cell behavior, however distantly. Although research examining the interaction between T cells and neural development is in its early stages, microglia have been shown to play an active role in neural circuit formation and patterning in utero and throughout life (Morimoto & Nakajima 2019). Mice genetically altered to lack T cells have shown differences in brain morphology (Rilett et al. 2015), suggesting that T cells may participate in shaping the developing brain, and vice versa.

Our results echo findings from recent single-cell sequencing studies in T cells from AD patients and aged individuals. We recently detected a cluster of CD8+ T cells with high expression of *KLRD1*, *KLRG1*, *GNLY*, *NKG7*, and several granzyme genes in immune cells isolated from surgically resected and post-mortem tissue (Olah et al. 2020). This is potentially consistent with the characterization of a clonally expanded population of CD8+ T cells within the cerebrospinal fluid of AD patients, whose expression of these genes correlates with their degree of clonality (Gate et al. 2020). CD8+ T cells from peripheral blood of AD patients have also shown levels of NKG7, GZMB, and GNLY expression that match natural killer cells (Xu & Jia 2021). Finally, another study of blood-derived T cells in supercentenarians suggests that this cytotoxic profile is not limited to disease. Hashimoto et al. (2019) found a cluster of cytotoxic CD4+ T cells in supercentenarians with many of these genes upregulated, especially in cells expressing late-differentiation markers. Our findings from co-expression network analysis confirm the trend of cytotoxic and senescent T cells in aging and AD. The appearance of many of these genes within our significant eQTL build upon previous research, by suggesting a contribution of genetic variants to these T cell phenotypes. Ascertaining the role of genetic variability behind age- or disease-related T cell cytotoxicity could lead to more personalized treatments for neurodegeneration and other conditions.

To that end, the detection of disease-associated eQTL in an aged and AD cohort warrants a closer look into the interindividual variability in T cell responses to neurodegeneration. Animal studies designed to examine T cell behavior in CNS disease, or test a T cell-targeted therapeutic strategy in a preclinical model, have yielded widely varying results for AD. Even within a T cell subtype such as regulatory T cells, different studies have found reduced (Baruch et al. 2015) or advanced (Baek et al. 2016) pathology upon depletion of Tregs. Th1-type CD4+ T cells specific for amyloid beta failed to lower amyloid levels in the hippocampus in one mouse model study (Ethell et al. 2006), while they improved amyloid clearance by microglia in another (Mittal et al. 2019), and amyloid-specific Th2 cells accomplished symptom reduction in AD mice even when introduced outside the CNS (Cao et al. 2009). While we know that T cell infiltration into the CNS occurs in AD (Merlini et al. 2018), more research is needed before we can better understand which T cell functions are helpful and hurtful in neurodegenerative disease. Precision medicine requires a deeper look into the source of genetic and environmental variability in T cell behavior, and T cell antigen specificity, during neurodegenerative disease.

Several recent eQTL studies have begun to detail key genes and pathways contributing to differential T cell phenotypes. Raj et al. (2014) conducted a large eQTL study of monocytes and naïve CD4+ T cells in young healthy subjects, finding that AD-associated variants among the cis-eQTL were mostly found in monocytes, while T cells cis-eQTL unsurprisingly held many of the autoimmune disease-related variants. Ye et al. (2014) focused mainly on transcript levels of cytokines from differentiated CD4+ T cells, but also conducted limited eQTL analysis. An extensive multi-omic study of several immune cell types, including T cells, by Chen et al. (2016) uncovered many variants related to autoimmune disease. Kasela et al. (2017) included CD8+ T cells as well as CD4+, and also detected autoimmune disease-related variants that affected the IRF1 and STAT1 signaling pathways. Bossini-Castillo et al. (2019) conducted a study using Tregs and focused especially on variants that could also be drug targets, finding several eQTL related to IL-10 signaling. Most recently, Nathan et al. (2021) used single-cell sequencing data from resting memory T cells to examine the interaction between eQTL and a wide variety of T cell states, an approach that will be vital in future studies, given the current expansion of single-cell sequencing technology.

These studies are an invaluable resource to mine genetic and epigenetic changes in T cells. However, their utility in elucidating how genetic variants may alter T cell behavior in age-related disease is limited, owing to the use of young, healthy subjects for sample collection. While our study is the first to use cells from aged and AD subjects to detect eQTL, it represents only a preliminary look into the complex network of T cell regulation in AD. Additional studies are warranted with a higher number of participants, to yield sufficient statistical power. We also recognize that the unique contribution of TEMRA cells to T cell phenotypes in neurodegeneration, as elaborated by Gate et al. 2020 and others, may have been obscured in our data, due to our sorting strategy. We plan to enrich the existing dataset with in vitro and in vivo studies to examine key genetic hits in AD models. With our planned follow-up experiments and others, we expect to make progress towards identifying causality in T cell genetic variants contributing to AD and other neurodegenerative diseases.

## Methods

### The Rush Religious Orders (ROS) Study and Rush Memory and Aging Project (MAP)

ROS, started in 1994, enrolls Catholic priests, nuns, and brothers, without known dementia, aged 53 or older from more than 40 groups in 15 states across the USA. Since January 1994, more than 1,450 participants completed their baseline evaluation, of whom 87% are non-Hispanic white, and the follow-up rate of survivors and autopsy rate among the deceased both exceed 90%. MAP, started in 1997, enrolls men and women without known dementia aged 55 or older from retirement communities, senior and subsidized housing, and individual homes across northeastern Illinois. Since October 1997, more than 2,200 participants completed their baseline evaluation, of which 87% were non-Hispanic white. The follow-up rate of survivors exceeds 90% and the autopsy rate exceeds 80%. Both studies were approved by an Institutional Review Board of Rush University Medical Center. All participants signed informed consent for detailed annual clinical evaluation, an Anatomic Gift Act, and a repository consent to allow their data and biospecimens to be share. All ROSMAP participants agree to organ donation at death. A subset of 450 ROS and all MAP participants agreed to annual blood draw. Cryopreserved PBMC were stored at baseline and annually from 2008 to present. More detailed descriptions of ROS, MAP, and CHAP can be found in prior publications (Bennett et al. 2018). ROSMAP resources can be requested at https://www.radc.rush.edu.

PBMC from 96 study autopsied participants were used in this study, of whom 48 had received a clinical diagnosis of AD dementia at the time of death, while 48 subjects without dementia served as controls. Brain tissue from each subject was analyzed post-mortem to detect pathological signs of neuritic plaques and neurofibrillary tangles.

### PBMC Isolation, Cell Sorting, and RNA Extraction

Peripheral blood mononuclear cells (PBMCs) were initially separated after collection in a yellow-top ACD tube, and later by Ficoll-Paque PLUS (GE Healthcare, Piscataway, NJ) gradient centrifugation, which markedly improved yield. PBMCs were frozen at a concentration of 1-3×10^7^ cells/ml in 10% DMSO (Sigma-Aldrich, St. Louis, MO)/90% FCS (Atlanta Biologicals, Lawrenceville, GA). After thawing, PBMCs were washed in 10 ml PBS. PBMCs were stained with monoclonal antibodies against human CD3, CD4, CD8, and CD45RO (BD Biosciences) in PBS plus 1% FCS. The following four T cell subsets were sorted by high-speed flow cytometry with a FACS Aria (BD Biosciences) to typically >98% purity in postsort analysis: CD3+CD4+CD45RO-, CD3+CD4+CD45RO+, CD3+CD8+CD45RO-, and CD3+CD8+CD45RO+. Total RNA was extracted using buffer TCL (Qiagen).

### Digital Gene Expression (DGE) 3’ *RNA-seq*

The RNA-seq libraries were prepared according to the Single Cell RNA Barcoding and Sequencing method originally developed for single cell RNA-seq (Soumillon et al., 2014) and adapted to extracted total RNA. Briefly, Poly(A)+ mRNA from extracted total RNA were converted to cDNA decorated with universal adapters, sample-specific barcodes, and unique molecular identifiers (UMIs) using a template-switching reverse transcriptase. Decorated cDNA from multiple samples were then pooled, amplified (10 PCR cycles), and prepared for multiplexed sequencing using a modified transposon-based fragmentation approach that enriched for 3′ ends and preserved strand information. RNA was sequenced using the Illumina HiSeq at the Broad Institute’s Broad Technology Labs and the Broad Genomics Platform using the High-throughput 3’ Digital Gene Expression (DGE) library (Soumillon et al., 2014).

### Gene Expression Quality Control and Differential Expression Analysis

Quality control on DGE was applied at sample and gene level. Genes with maximum cell count value of at least three and non-zero values in over twenty percent of samples were included in differential expression analysis. Expression values were normalized to counts per million (CPM). We correlated gene expression with AD and related neuropathological traits including a) clinical diagnosis of dementia, b) autopsy confirmed diagnosis of AD, c) global pathology defined as global measure of pathology based on the scaled scores of five brain regions, d) Braak Stage (Braak & Braak 1991), e) PHFtau tangle density across eight brain regions, and f) area occupied by β-amyloid across eight brain regions. The EdgeR package in R (Robinson et al. 2010) was used to conduct differential expression. Voom transformation (Law et al. 2014) was applied to gene expression data and regression analysis was used to test association with the AD and neuropathological traits. Models were adjusted for age at death, sex, and total cell counts for each sample. Significant genes were determined at FDR≤0.05.

### Genotyping and Imputation

Subjects from the ROS and MAP cohorts imputed dosage values were used in our study. In these cohorts, DNA was extracted from whole blood or frozen post-mortem brain tissue. Genotype data was generated using the Affymetrix Genechip 6.0 platform at the Broad Institute’s Genetic Analysis Platform or the Translational Genomics Research Institute. Both sets of data underwent the same quality control (QC) analyses in parallel using the PLINK toolkit (http://pngu.mgh.harvard.edu/~purcell/plink/) and quality controlled genotypes were pooled. The QC process included a principal components analysis using default parameters in EIGENSTRAT (Price et al. 2006) to identify and remove population outliers. Imputation in ROS and MAP was performed using MACH software (version 1.0.16a) and HapMap release 22 CEU (build 36).

### Gene co-Expression Module Assembly

To identify modules of co-expressed genes and their associations with cell types and traits, we applied weighted gene co-expression network analysis (WGCNA, Langfelder & Horvath 2008) to the sorted cell RNA-seq data. Sample expression values were normalized first by randomly subsampling to 10,000 transcripts, followed by normalization to counts per million (CPM). These normalized values were then used as input to WGCNA, using the top 3000 genes with highest % CV and the following WGCNA parameters: power = 6, TOMtype = “unsigned”, minModuleSize=30, and mergeCutHeight=0.25. This yielded three modules (apart from the “grey” or unassigned gene module), whose eigengenes were then used for association analyses with cell classes and disease traits.

### Expression Quantitative Trait Loci (eQTL) Testing

We computed eQTLs with common single nucleotide polymorphisms (SNPs) (minor allele frequency > 0.1). First, we identified independent SNPs across the genome by pruning for LD (R^2^ <0.2 between all pairs of SNPs). Then, we used the MatrixeQTL (Shabalin 2012) to test for associations of SNPs with gene expression in each T-cell subtype independently. Association tests were adjusted for age at death, sex, and clinical AD status. Significant pairings (eQTLs) were classified as *cis* (SNPs within 1 Mb of the gene) or *trans* (SNPs at least 1 Mb away from the transcription start site or on a different chromosome). Significant eQTLs were determined at a false discovery rate≤0.05, calculated independently for the cis and trans-eQTLs.

### Venn Diagrams and Manhattan plots for trans eQTLs

Venn diagrams were assembled using the VennDiagram package in R (Chen & Boutros 2011) with lists of unique genes and unique SNPs from each of the four T cell subsets. The qqman package (Turner 2014) was used to create Manhattan plots of SNPs from the trans-eQTL. SNPs with a p-value under 1 x 10^−14^ were annotated manually on the Manhattan plot.

### GWAS Catalog Analysis

We downloaded the GWAS catalog v1.0.2 with annotation for SNPs and genotyping arrays. The catalog was filtered to retain studies of traits falling into four categories: a) immune, b) autoimmune, c) neurodegenerative, and d) neuropsychiatric. Significant cis- and trans-eQTLs were cross-referenced with GWAS catalog entries based on their dbSNP IDs. Genes that harbored eQTLs or whose expression was influenced by genetic variation in the four disease categories were further prioritized for pathway analysis.

### Pathway Analysis

Pathway analysis was done using the enrichR package in RStudio (Chen et al. 2013, Kuleshov et al. 2016, Xie et al. 2021). Unranked gene lists from the co-expression modules or from trans-eSNPs in coding regions were used as input. GO Biological Processes 2021 was used as the database (Ashburner et al. 2000, Gene Ontology Consortium 2020). The enrichr function was used to calculate the most highly enriched GO pathways represented by the input genes. Dot plots were generated from the enrichr results using the PlotEnrich function and modified with the ggplot2 package.

### Statistical Methods

Multiple testing correction for differential gene expression, eQTL detection, and pathway analysis was done by calculating the false discovery rate and reporting results with a q-value less than 0.05. When comparing gene expression with AD neuropathological traits in figure 1c, results were reported as nominal p-values with no multiple testing correction.

## Supplementary

<u>Supplementary Table 1: Raw data from RNA-sequencing.</u> Count matrix from 3’ digital gene expression (DGE) from 4 sorted T cell subsets in 96 ROSMAP participants. Libraries were prepared according to the Single Cell RNA Barcoding and Sequencing method originally developed for single cell RNA-seq. Sequencing was done using the Illumina HiSeq platform. Further details can be found in the Methods section.

<u>Supplementary Table 2: Association of gene expression with AD traits.</u> Listed here are the genes significantly associated with one or more AD pathological traits. Pathological trait data is collected from ROSMAP participants, gene expression data is our own. The values below are reported for each of 6 pathological measurements: pathoAD (the presence or absence of a neuropathological diagnosis of AD), clinAD (the presence or absence of a clinical diagnosis of AD), gpath (a measure of global AD pathology), Braak (a measure of neurofibrillary tangle pathology in terms of Braak staging), amyloid (a measure of amyloid plaque burden), and tangles (a measure of tau tangle burden).

<u>Supplementary Table 3: Genes in three WCGNA modules.</u> Listing of genes in three modules obtained from Weighted Gene Co-Expression Network Analysis (WGCNA). The WGCNA package in R (developed by Langfelder & Horvath) was used with RNA-sequencing data as input, with sample expression values normalized. For further details, see the Methods section.

<u>Supplementary Table 4: Cis-eQTL organized by T cell subtype.</u> Full list of genes and SNPs in cis-eQTL after use of the Matrix eQTL method, with multiple testing correction applied to limit results to those with a false discovery rate (FDR) under 0.05. Results organized by T cell subset. Further details about Matrix eQTL, and eQTL generation, can be found in Methods.

<u>Supplementary Table 5: Trans-eQTL organized by T cell subtype.</u> Full list of genes and SNPs in trans-eQTL after use of the Matrix eQTL method, with multiple testing correction applied to limit results to those with a false discovery rate (FDR) under 0.05. Results organized by T cell subset. Further details about Matrix eQTL, and eQTL generation, can be found in Methods.

<u>Supplementary Table 6: Trans-eQTL with SNPs found in the GWAS catalog.</u> Annotation of SNPs from trans-eQTL which are also found in the GWAS catalog. Includes information about chromosomal location of the SNP, the associated GWAS trait, and further metrics of the study detecting the SNP-trait correlation. All associations are from GWAS catalog v1.0.2.

<u>Supplementary Table 7: List of traits used for selected disease association categories, and trans-eQTL with SNPs found in those categories.</u> Listing of all phenotypes found in the GWAS catalog, followed by the four categories used to create the stacked bar graph and Circos plots in figure 4. Trans-eQTL whose SNP is associated with one of the phenotypes in these four categories are found in the other tabs of this spreadsheet, organized by cell type.

<u>Supplementary Figure 1: Pathway analysis for genes represented by the trans-eQTL.</u> Top 20 most highly enriched GO pathways from EnrichR results for a) CD4+CD45RO-, b) CD4+CD45RO+, c) CD8+CD45RO-, d) CD8+CD45RO+. Dot size represents number of differentially expressed genes from the trans-eQTL, x axis location represents ratio of a GO pathway represented by trans eQTL genes, dot color represents P value.

## Acknowledgments

We thank the participants in the Religious Orders Study and the Rush Memory and Aging Project for donating their data and biospecimen. This work was supported by the US National Institutes of Health grants R01AG067581 and R01NS124771 and the Ludwig Family Foundation (to W.E) and the NIH R01AG043617 to E.M.B., and P30AG10161, P30AG72975, R01AG15819, R01AG17917, and U01AG61356 to D.A.B.. D.D. is a recipient of a TL1 Precision Medicine Predoctoral Fellowship (TL1TR001875).

## Authors Contributions

W.E. and E.M.B. designed and implemented the study. T.B. and M.C. conducted cell sorting experiments. D.A.B. contributed the human blood samples and clinical and neuropathological data and critically reviewed the paper. D.D., V.M., E.M.B., B.V., and W.E. performed statistical analyses and interpretation of results. D.D., B.V., and W.E. wrote the manuscript.

