## Supplementary figures and images for "Genotype-Phenotype Correlation of T Cell Subtypes Reveals Senescent and Cytotoxic Genes in Alzheimer’s Disease"

### Supplementary figure 1

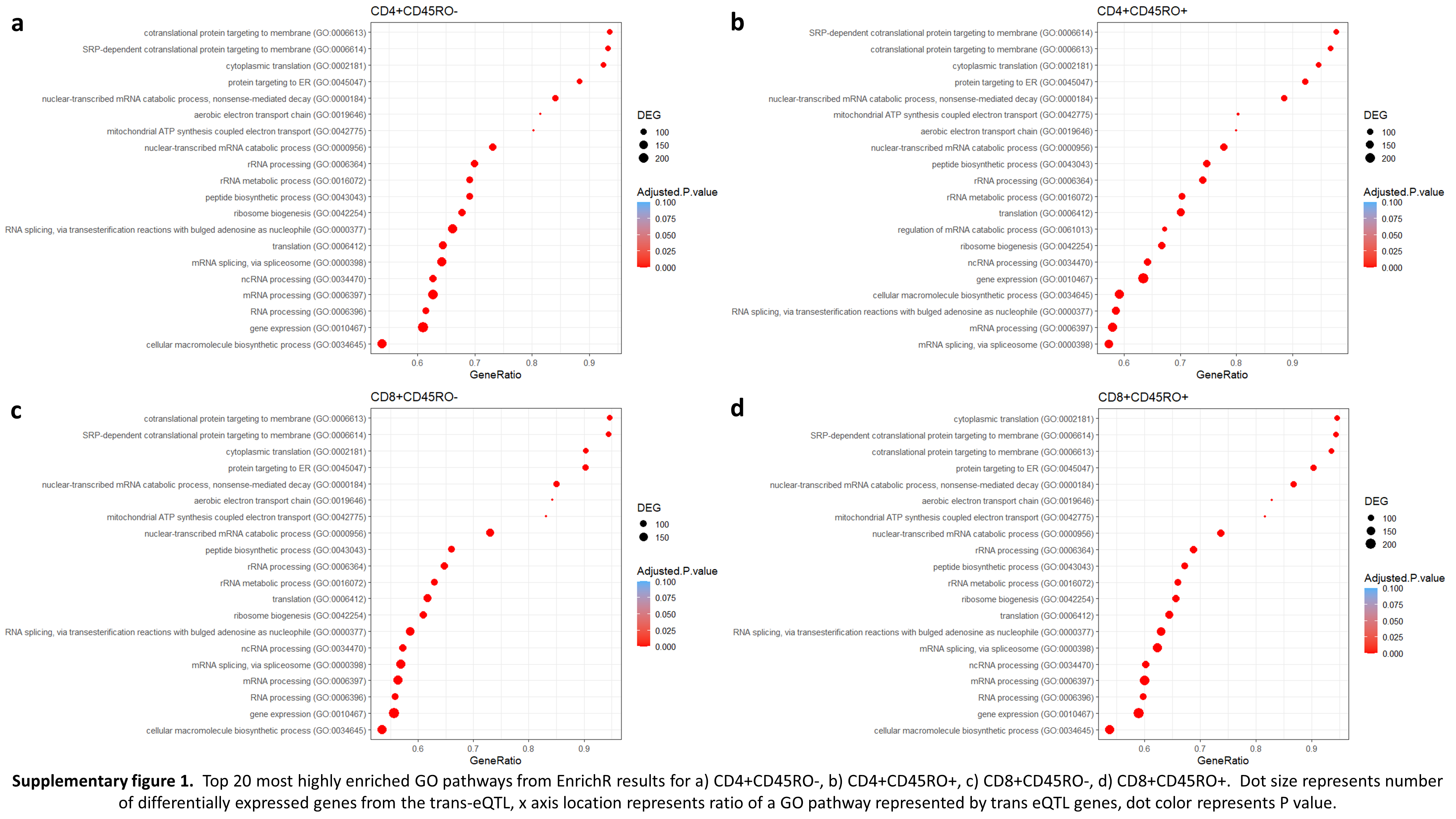
